# Root endophyte sulfur metabolites enhance redox balance and drought tolerance in Arabidopsis

**DOI:** 10.64898/2026.06.15.732246

**Authors:** Büsra Elkatmis, Rabeaa S. Alkhateeb, Catherine Mannes, Rewaa Jalal, Marilia Trapp, Philipp Westhoff, Ceyda Ozkan, Giulia Marie Thelen, Baoda Han, Maged M. Saad, Stanislav Kopriva, Heribert Hirt

**Author notes:** **Corresponding Authors**: Stanislav Kopriva, Heribert Hirt, **Email:**.

## Abstract

Drought is a major consequence of climate change and significantly limits crop productivity. Plant growth-promoting bacteria offer a promising solution to mitigate drought stress. The root endophyte *Pseudomonas argentinensis* SA190 has been shown to enhance plant performance under drought stress conditions, but the mechanistic basis of SA190’s beneficial effects remained unclear. Given the pivotal role of sulfur-containing compounds in abiotic stress responses, we investigated multiple sulfur-related Arabidopsis mutants under drought stress. We found that SA190 enhances sulfate uptake and promotes glutathione (GSH) accumulation in shoots under stress conditions. SA190 treatment improved the GSH/GSSG ratio, indicating an enhanced redox balance under drought. Selective inhibition of Arabidopsis GSH biosynthesis using buthionine sulfoximine (BSO) confirmed the essential contribution of bacterial GSH to drought stress. In addition, by generation and use of bacterial mutants deficient in the GSH synthesis pathway, we show that the bacteria directly provide Arabidopsis with either GSH or it’s precursor γ-EC. In summary, SA190 promotes drought tolerance by supplying the host plant with additional GSH thereby maintaining cellular redox homeostasis and enhancing drought stress resilience.

## Introduction

Drought is one of the most severe abiotic stresses limiting plant growth and agricultural productivity worldwide (1, 2). It reduces water availability, impairs photosynthesis, nutrient uptake, and cellular metabolism, and leads to the accumulation of reactive oxygen species (ROS) (3, 4). Excessive ROS disrupt membrane integrity, damage proteins, and impair essential cellular functions, ultimately resulting in reduced biomass and yield (5). To counteract drought-induced oxidative damage, plants activate a range of defense mechanisms, including enzymatic antioxidants such as catalase (CAT), superoxide dismutase (SOD), glutathione reductase (GR), and glutathione S-transferase (GST), along with non-enzymatic osmolytes and/or antioxidants like proline, glycine betaine, carotenoids, and glutathione (GSH) (6, 7). Among these, sulfur and sulfur-containing compounds play a pivotal role in abiotic stress tolerance (8, 9). In particular, GSH, a central sulfur-containing tripeptide, is essential for ROS detoxification, redox homeostasis, and cellular adaptation under stress conditions (10). Multiple studies demonstrated the importance of GSH in protection against oxidative stress across diverse species (11, 12). For instance, exogenous GSH treatment mitigated oxidative stress in wheat seedlings exposed to heat and drought, enhancing shoot growth traits, such as plant height and branch number (11). Similarly, in tomato, GSH application reduced H_2_O_2_ accumulation under salt stress, increased the GSH/GSSG ratio, and enhanced activities of antioxidant enzymes such as SOD and CAT, thereby improving redox balance (12).

In addition to intrinsic antioxidant defenses, plants can benefit from interactions with beneficial microbes to enhance stress tolerance. Plant growth-promoting bacteria (PGPB) play a key role in improving plant performance under stress by modulating key physiological processes, including phytohormone regulation, nutrient acquisition, and hydrolytic enzyme production (13, 14). For example, *Piriformospora indica* colonization improves drought tolerance in maize by enhancing root architecture, activating antioxidant pathways, and reprogramming gene expression, especially in sulfur and carbon metabolism (15). Similarly, *Enterobacter* sp. SA187 enhances growth in *Arabidopsis thaliana* under salt stress by supplying 2-keto-4-methylthiobutyric acid (KMBA), a sulfur-containing ethylene precursor that contributes to stress mitigation (16). In addition to these, *Pseudomonas argentinensis* SA190, a desert adapted bacterium, has been shown to enhance plant biomass and drought resilience in Arabidopsis (17, 18). However, the mechanistic basis of this beneficial interaction, particularly its connection to sulfur metabolism and redox regulation, remains poorly understood.

Here, we investigate how SA190 modulates host sulfur metabolism under 25% polyethylene glycol (PEG)-induced drought stress. We show that SA190 enhances GSH accumulation even when the plant’s own biosynthetic capacity is impaired, indicating that the bacteria may contribute to the plant’s GSH pool or by modulating the plant’s pathways. SA190 also enhances redox balance, reduces superoxide accumulation, and activates metabolites involved in GSH biosynthesis. Moreover, we demonstrate that bacterial glutathione biosynthesis, specifically through the *gshA* gene, is critical for root colonization and bacterial fitness. These findings reveal a previously uncharacterized role for microbial sulfur metabolism in promoting plant drought tolerance and position SA190 as a promising candidate for improving crop resilience under drought stress conditions.

## Results

### SA190 Enhances Biomass of Sulfur Mutants, Sulfate Uptake, and Glutathione Levels in Arabidopsis

To assess the role of sulfur metabolism in the SA190-mediated protection against PEG-induced drought stress, we evaluated the *sultr1;2, apr2*, and *cad2-1* mutants. Under control conditions, *sultr1;2* and *cad2-1* mutants exhibited slightly reduced biomass relative to *Arabidopsis thaliana* Col-0 (Figure 1A), reflecting impairments in sulfate uptake (*sultr1;2*) (19–21) and glutathione biosynthesis (*cad2-1*) (22). Under PEG-induced drought stress, both *sultr1;2* and *cad2-1* mutants displayed heightened drought sensitivity, with pronounced reductions in biomass (Figure 1B). The *apr2* mutant, defective in sulfate reduction (23), also showed reduced biomass under 25% PEG-induced drought stress. The phenotypes of the mutants were consistent with the biomass results (Supplementary Figure 1 and 2). Treatment with SA190 significantly enhanced biomass in all three mutants, particularly under PEG-induced drought stress. In *sultr1;2*, SA190 led to a 3-fold increase in biomass under control conditions and 35-fold increase under PEG-induced stress compared to mock treatment. Similarly, under PEG-induced drought stress SA190 promoted a 5-fold and 25-fold increase in biomass in the *apr2* and the *cad2-1* mutants, respectively. These findings show that glutathione, and possibly other sulfur-containing compounds, are crucial for the response of Arabidopsis to PEG-induced drought stress. They also suggest that SA190 may support plant survival under these conditions by promoting uptake or biosynthesis of sulfur-containing compounds.

**Figure 1.**
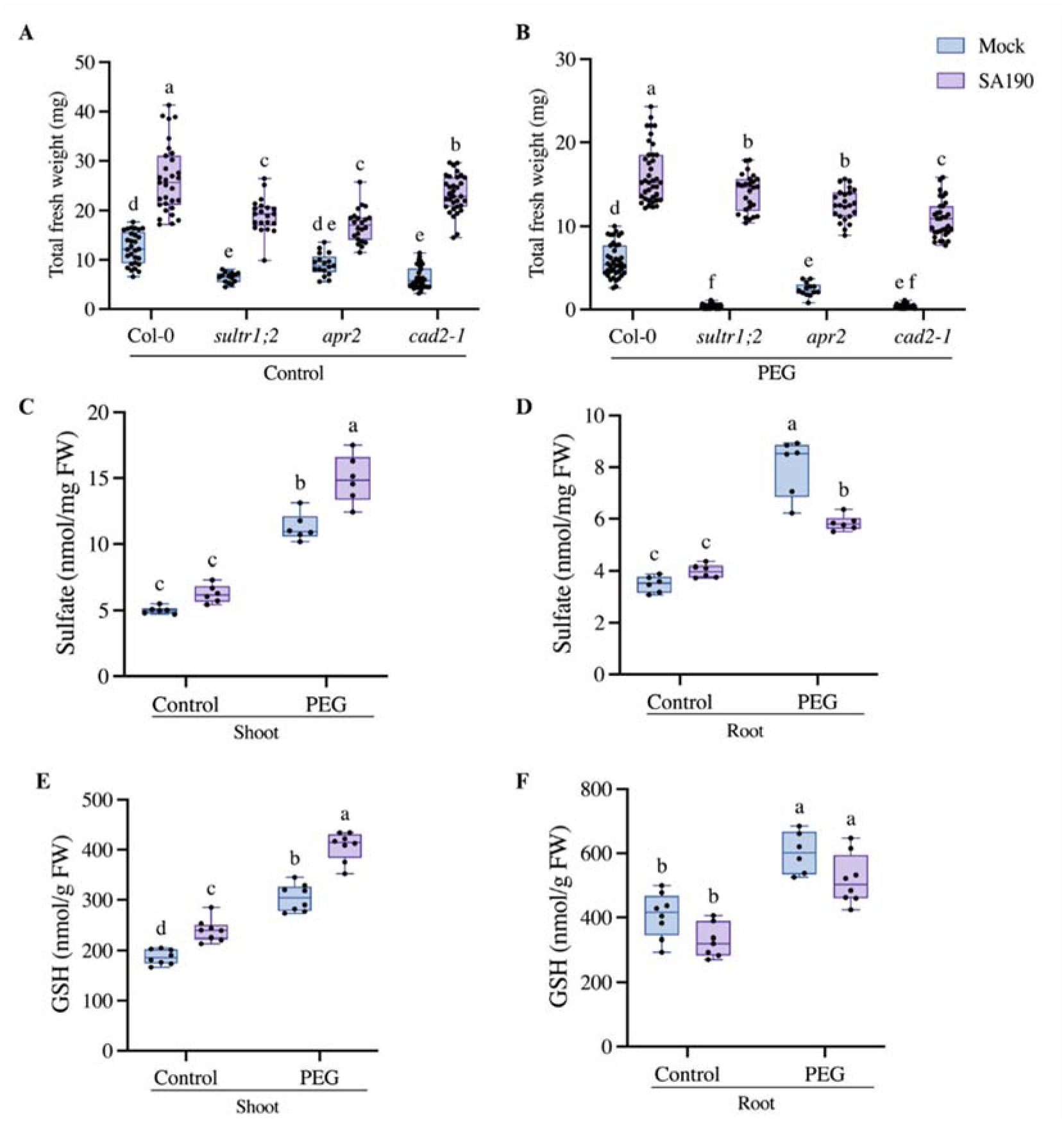
Effect of *Pseudomonas argentinensis* SA190 on sulfur-deficient Arabidopsis mutants, sulfate uptake, and glutathione (GSH) levels under control (non-stress) and 25% PEG-induced drought stress conditions. Total fresh weight of non-colonized (mock) and SA190-colonized *Arabidopsis thaliana* Col-0 grown on control (A) or 25% PEG-induced drought stress (B) for 15 days. Sulfate content in shoot (C) and root (D) samples under control and 25% PEG-induced drought stress conditions. GSH levels in shoot (E) and root (F) tissues of mock- and SA190-inoculated plants under the same conditions. The plant assay was performed in three biological replicates and two technical replicates (n=36). Whiskers represent minimum and maximum values, the horizontal line indicates the median. Different letters indicate statistically significant differences based on two-way ANOVA (p < 0.05).

To determine whether SA190 influences the accumulation of sulfur-containing compounds in plants, we measured sulfate and glutathione (GSH) levels in *Arabidopsis thaliana* Col-0 under control and 25% PEG-induced drought stress conditions. Sulfate and GSH levels were significantly elevated in both shoot and root under PEG-induced drought stress compared to control condition, highlighting the potential role of sulfur metabolism in the drought response (Figure 1). While under control condition, SA190 treatment did not affect sulfate concentration, under PEG-induced drought stress sulfate levels were significantly increased in the shoot of SA190 treated plants relative to mock (Figure 1C). In contrast, sulfate levels in the root of PEG-treated plants were significantly reduced with SA190 treatment (Figure 1D), suggesting enhanced translocation of sulfate from the root to the shoot potentially to support increased sulfur demand in aerial tissues.

Sulfate levels in shoots of the sulfur-related mutants (*sultr1;2, apr2, cad2-1*) showed that *sultr1;2* had reduced sulfate levels compared to Col-0 under control and PEG stress conditions, and SA190 did not restore its accumulation in *sultr1;2* (Supplementary Figure 3A and B). Although SA190 rescued the *sultr1;2* growth phenotype under PEG stress, sulfate levels in shoots remained low. A similar pattern was also observed in roots (Supplementary Figure 3C and D). The *apr2* mutant had higher shoot sulfate accumulation than Col-0 under control and PEG stress conditions due to lower rate of sulfate reduction, and SA190 further increased it. In contrast, *cad2-1* showed reduced shoot sulfate compared to Col-0 under control and PEG stress, however, SA190 enhanced the sulfate level in *cad2-1*. Analysis of sulfate levels in the mutants suggests that SA190 may provide a sulfur-containing compound functioning downstream of the sulfur assimilation pathway.

Consistently, in the wild type, SA190 significantly increased GSH levels in the shoot compared to mock under both control and stress conditions (Figure 1E), whereas, GSH in the root remained unchanged (Figure 1F). These results suggest that to mitigate drought-induced oxidative stress, SA190 promotes shoot-specific GSH accumulation, potentially through enhanced sulfate uptake and transport in the root, followed by preferential GSH synthesis in the shoot. Alternatively, SA190 might provide GSH to the plants directly.

### SA190 Enhances GSH Accumulation in Plants Deficient in Glutathione Biosynthesis

In order to test whether SA190 contributes directly to the GSH pool in plants, we first assessed if SA190 might compensate a buthionine sulfoximine (BSO) inhibition of GSH synthesis in Arabidopsis. BSO is a specific inhibitor of γ-glutamylcysteine synthetase (γ-ECS), the rate-limiting enzyme in GSH biosynthesis, and thereby suppresses endogenous GSH production in plants (24). Treatment with 1 mM BSO markedly inhibited Arabidopsis growth under both control and 25% PEG-induced drought stress conditions, resulting in visible phenotypic alterations (Supplementary Figures 4 and 5). Primary root length was significantly reduced in BSO-treated plants compared to control plants under both conditions (Figure 2A and B), which has in turn contributed to a significant reduction in plant biomass (Figure 2C and D). On the other hand, BSO did not affect growth of SA190 (Supplementary Figure 6). The exogenous application of 0.5 mM GSH (Figure 2C and D) enhanced Arabidopsis survival under 25% PEG-induced drougt stress, but had no effect under control conditions, indicating that GSH alone can improve plant survival under stress.

**Figure 2.**
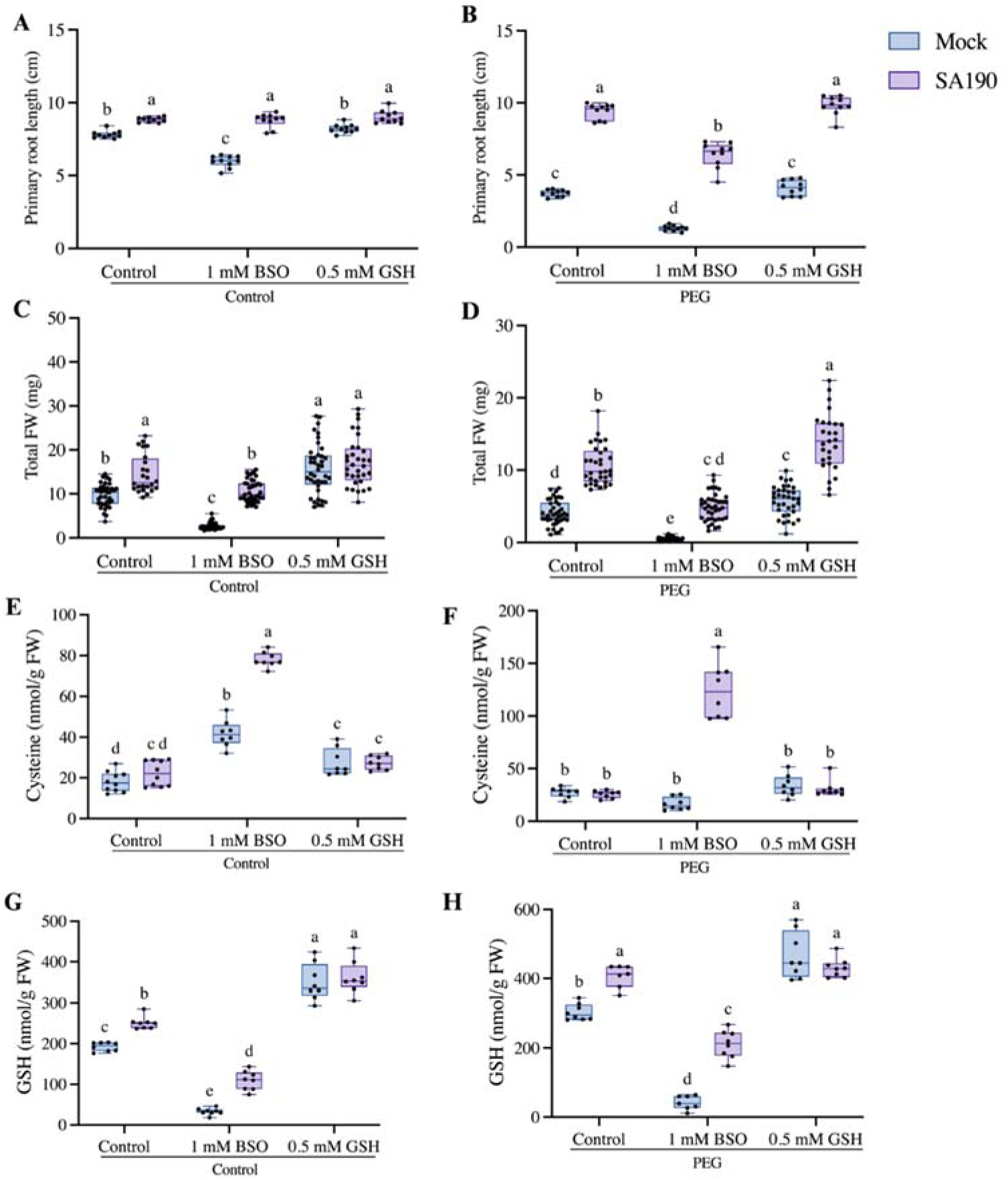
Effect of *Pseudomonas argentinensis* SA190 on Arabidopsis growth in the presence of BSO and GSH under control (non-stress) and 25% PEG-induced drought stress conditions. Primary root length of non-colonized and SA190-colonized *Arabidopsis thaliana* Col-0 under control, 1 mM BSO, and 0.5 mM GSH treatments on control (A) and 25% PEG-induced drought stress (B). Root length was measured using ImageJ (version 1.53t). Total fresh weight of non-colonized and SA190-colonized plants under the same treatments on control (C) and 25% PEG-induced drought stress (D) conditions. Cysteine levels in shoot tissue under the same treatments on control (E) and 25% PEG-induced drought stress (F). Glutathione (GSH) levels in shoot tissue under the same treatments on control (G) and 25% PEG-induced drought stress (H). The plant assay was performed in three biological replicates and two technical replicates (n=36). Whiskers represent minimum and maximum values, the horizontal line indicates the median. Different letters denote statistically significant differences based on two-way ANOVA (p < 0.05).

Inoculation with SA190 alleviated the negative effects of BSO treatment. Compared to mock, plants inoculated with SA190 showed increased primary root length and higher biomass accumulation on BSO containing agar plates under both control and stress conditions. Exogenous GSH supplementation coupled with SA190 treatment resulted in an additive beneficial effect, as the plants treated with both, GSH and SA190, showed the highest biomass accumulation under stress (Figure 2D). Improved plant growth therefore suggests a synergistic effect between bacterial presence and GSH supplementation.

We then quantified cysteine and GSH in the presence and absence of SA190. In control plants, colonization with SA190 under control as well PEG-induced drought stress conditions did not affect cysteine levels (Figure 2E and F). However, GSH levels were significantly enhanced by the colonization with the bacteria (Figure 2G and H), indicating that SA190 affects GSH synthesis in the host plant. Under BSO treatment, non-colonized shoot samples accumulated cysteine under control conditions (Figure 2E) and exhibited highly diminished GSH levels (Figure 2G and H) compared to control. These observations confirmed the effective inhibition of GSH biosynthesis upon BSO treatment. SA190 inoculation further increased cysteine levels in BSO-treated plants under both conditions (Figure 2E and F). Importantly, under both non-stressed and stressed conditions, SA190 treated plants showed increased GSH levels compared to non-colonized shoot samples (Figure 2G and H). Exogenous GSH supplementation restored shoot GSH content under both control and PEG-induced drought stress conditions irrespective of SA190 presence (Figure 2G and H). These results suggest that SA190 may contribute to GSH accumulation under PEG-induced drought stress, by supplying GSH to the plant.

To further examine whether SA190 may supply glutathione to the host plant, we analyzed the Arabidopsis glutathione-deficient mutant *cad2-1* in the presence of SA190. Under PEG-induced drought stress, *cad2-1* mutants exhibited a marked reduction in GSH levels compared to Col-0, confirming their compromised glutathione biosynthetic capacity. Notably, SA190 treatment significantly increased GSH levels in *cad2-1* under PEG-induced drought stress conditions (Supplementary Figure 7). These findings suggest that SA190 may contribute to the host glutathione pool, supporting the hypothesis that the bacteria may supply glutathione to the plant.

Altogether, these findings demonstrate a central role for SA190 in enhancing sulfur assimilation and promoting glutathione accumulation under 25% PEG-induced drought stress. In particular, the ability of SA190 to enhance GSH accumulation even when GSH biosynthesis is compromised supports a mode of beneficial interaction in which the bacteria facilitate sulfur metabolism and contribute to improved drought tolerance in Arabidopsis.

### SA190 Enhances Redox Homeostasis and Reduces Superoxide Accumulation Under Drought Stress

Glutathione (GSH) is a major antioxidant that plays a vital role in maintaining cellular redox homeostasis (25). Alongside its reduced form (GSH), the oxidized form, glutathione disulfide (GSSG), contributes as well to redox regulation (25, 26). Lower GSH/GSSG ratios serve as a key indicator of oxidative stress and redox imbalance in plants (26).

In order to assess the impact of SA190 on redox status within the plant, we quantified GSSG levels and calculated the GSH/GSSG ratio in Arabidopsis in response to SA190 treatment. Under 25% PEG-induced drought stress, GSSG levels significantly increased in the plant shoots compared to control conditions (Figure 3A), indicating the presence of oxidative stress. SA190 treatment markedly reduced GSSG levels in the shoot under PEG-induced drought stress, suggesting a mitigation of oxidative damage (Figure 3A). Consistently, SA190 significantly enhanced the GSH/GSSG ratio in the shoot under PEG-induced drought stress conditions, indicating improved antioxidant capacity and restored redox homeostasis (Figure 3B). In contrast, the GSSG level and the GSH/GSSG ratio in roots remained largely unchanged by SA190 (Figure 3C and 3D), potentially reflecting lower oxidative pressure or tissue-specific differences in glutathione metabolism.

**Figure 3.**
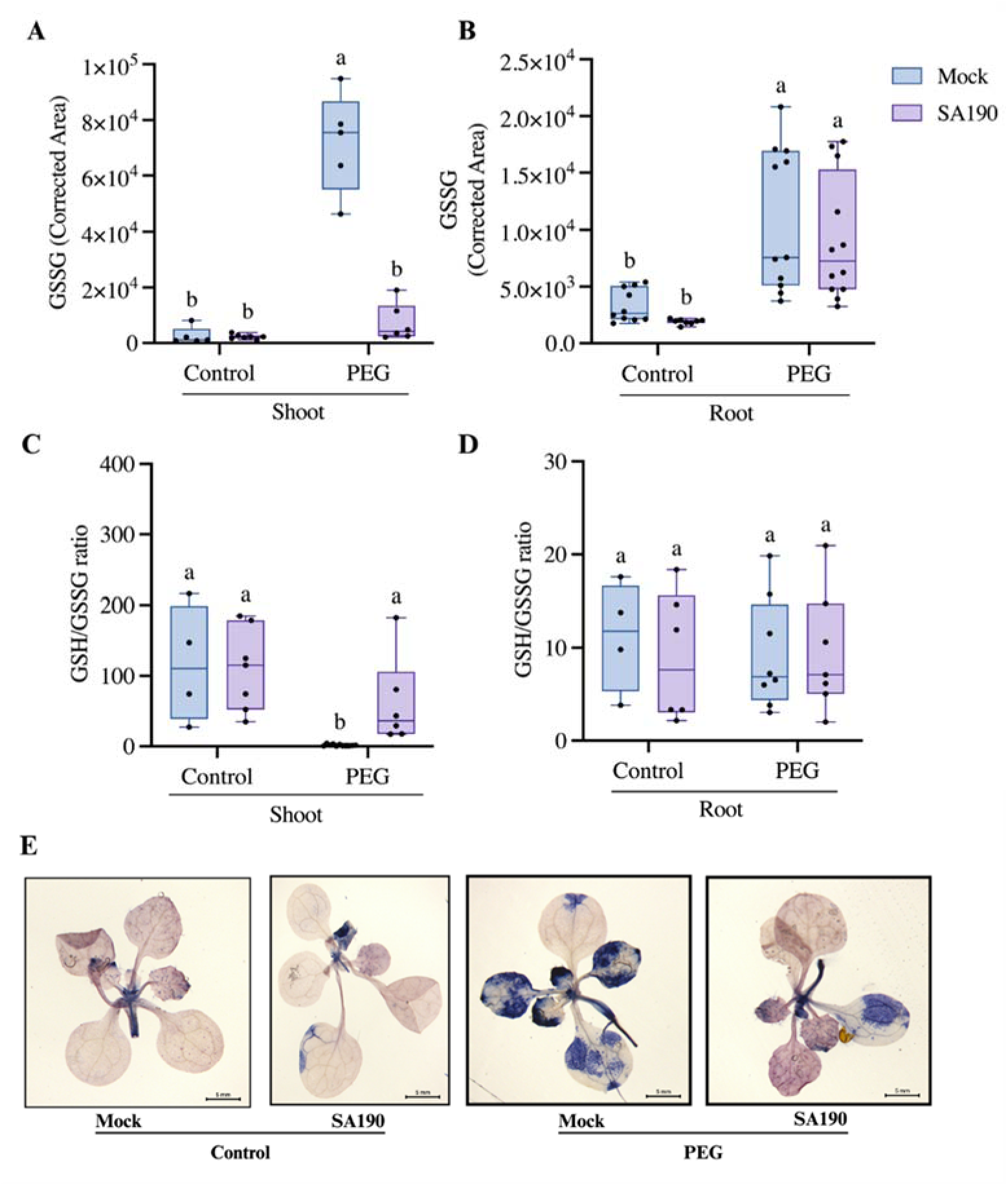
Effect of *Pseudomonas argentinensis* SA190 on redox homeostasis under control (non-stress) and 25% PEG-induced drought stress conditions. The GSSG level of non-colonized and SA190-colonized *Arabidopsis thaliana* Col-0 under control and 25% PEG-induced drought stress conditions for the shoot (A), the GSH/GSSG ratio of non-colonized and SA190-colonized *Arabidopsis thaliana* Col-0 under control and 25% PEG stress conditions for the shoot (B), the GSSG level of non-colonized and SA190-colonized *Arabidopsis thaliana* Col-0 under control and 25% PEG-induced drought stress conditions for the root (C), The GSH/GSSG ratio of non-colonized and SA190-colonized *Arabidopsis thaliana* Col-0 under control and 25% PEG stress conditions for the root (D). In the boxplot, the whiskers represent the minimum and maximum values, while the horizontal line represents the median. Different letters indicate statistical differences based on two-way ANOVA (p < 0.05). Effect of SA190 on superoxide accumulation in 7-day-old seedlings under control and 25% PEG-induced drought stress asessed by nitro blue tetrazolium (NBT) staining (F).

To further evaluate the impact of SA190 on reactive oxygen species (ROS) levels, we performed nitroblue tetrazolium (NBT) staining in SA190-colonized and non-colonized seedlings (Figure 3E). Strong superoxide accumulation marked by the extent of purple stained area was evident in the shoots of non-colonized seedlings under PEG-induced drought stress. In contrast, SA190 visibly reduced superoxide levels, demonstrating that SA190 attenuates ROS accumulation and alleviates oxidative stress in seedlings under PEG-induced drought stress. On the other hand, no differences were observed between the treatments under control conditions (Figure 3E).

### SA190 Activates the Host Metabolism Involved in Glutathione (GSH) Biosynthesis

To investigate the impact of SA190 on the sulfur metabolism and glutathione (GSH) biosynthesis of the host, we performed metabolic profiling of colonized Arabidopsis shoot and root tissues under control and 25% PEG-induced drought stress conditions. In the absence of SA190, plants subject to drought stress showed a distinct change in metabolic profiles compared to non-stressed plants (Figure 4A and B). As illustrated in the heatmaps, under 25% PEG-induced stress, the levels of myo-inositol, 5-oxoproline, aspartate, proline, and glycine were significantly elevated in root and shoot samples compared with control conditions (Figure 4A and B). These metabolites are associated with osmoprotection, nitrogen storage, and redox buffering (27–30), thus their presence is attributed to the physiological response during adaptation to the negative effects of drought. Notably, treatment with SA190 suppressed the accumulation of aspartate, proline, and glycine in shoot tissues under PEG-induced drought stress condition, suggesting that the presence of SA190 may modulate plant stress metabolism. On the other hand, SA190 treatment significantly increased the abundance of serine, methionine, asparagine, glutamine, and glutamate under PEG-induced drought stress compared to mock treatment (Figure 4A). Metabolites involved in plant sulfur metabolism and glutathione (GSH) biosynthesis are shown in Figure 4C. Among the up-regulated metabolites was glutamate, which plays a central role as an intermediate in amino acids biosynthetic pathways by serving as a precursor for the biosynthesis of glutamine, proline, methionine, and aspartate (31). Another up-regulated metabolite was serine which indirectly contributes to GSH biosynthesis by supplying O-acetylserine (OAS), an intermediate to cysteine biosynthesis (32). These coordinated metabolic changes indicate that SA190 may enhance GSH biosynthesis by promoting both sulfur and nitrogen assimilation. Together, our findings suggest that SA190-induced metabolic reprogramming under PEG-induced drought stress enhances the plant’s antioxidant capacity and stress resilience through targeted regulation of GSH-associated pathways.

**Figure 4.**
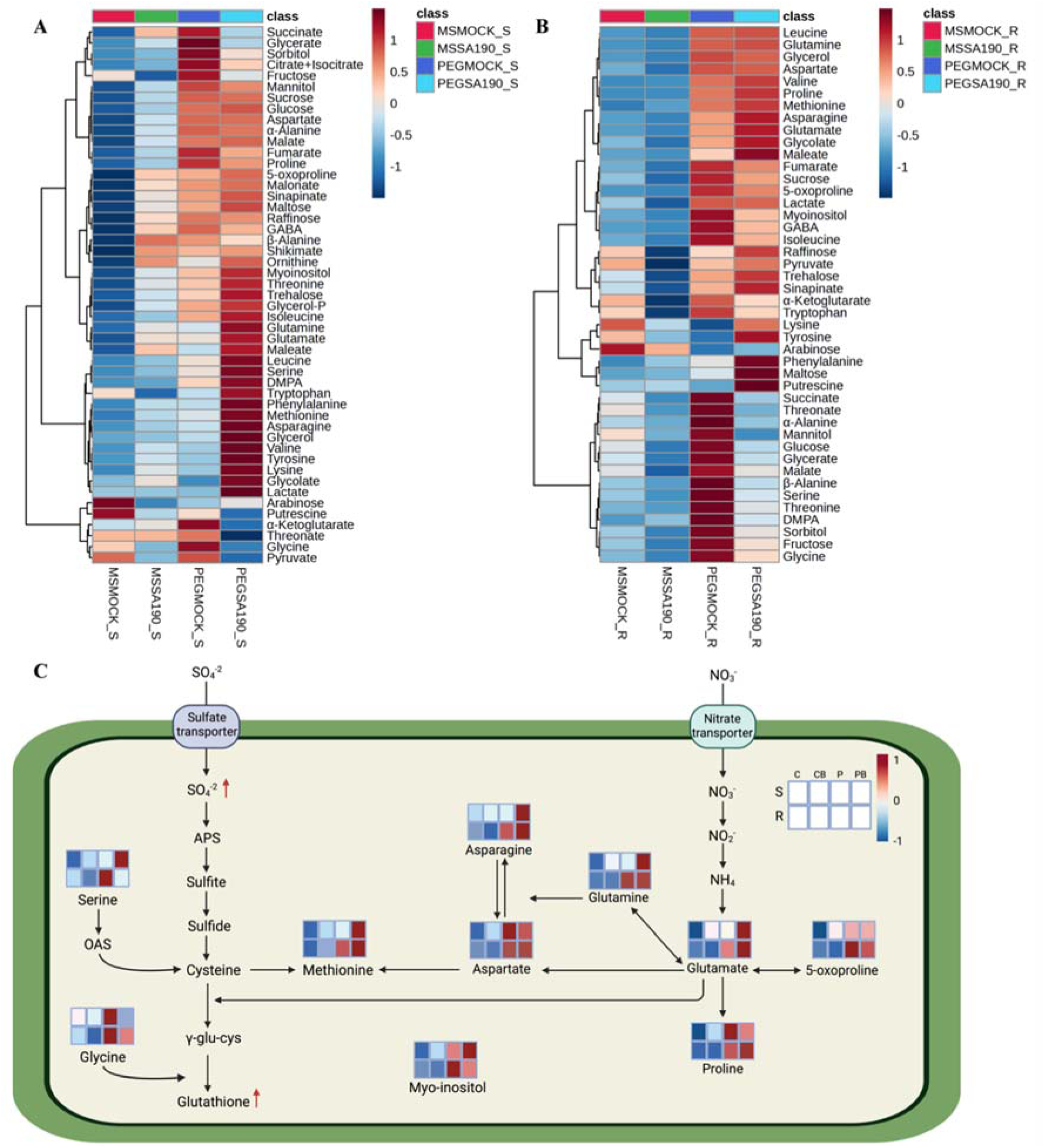
*Pseudomonas argentinensis* SA190 modulates sulfur-related metabolites and glutathione biosynthesis pathways under control (non-stress) and 25% PEG-induced drought stress. Heatmap of metabolites in shoot tissues of *Arabidopsis thaliana* treated with mock or SA190 under control (non-stress) and 25% PEG-induced drought stress conditions (A). Heatmap of metabolites in root tissues under the same conditions (B). Metabolic pathway map illustrating the connection between SA190-responsive metabolites involved in sulfur-related and glutathione (GSH) biosynthesis (C). S, shoot; R, root; C, Control (non-stress); CB, bacterial treatment under control condition (non-stress); P, PEG stress; PB, bacterial treatment under PEG stress. Metabolite levels are color-coded based on statistical significance: red indicates significant up-regulation; blue indicates significant down-regulation. Metabolic pathway map was created using Biorender.com (https://BioRender.com/qr8pqwq).

### Microbial gshA involved in Glutathione Biosynthesis Plays a Critical Role in Plant Survival under Drought Stress

The effects of SA190 on Arabidopsis GSH levels can be explained by either stimulation of sulfate assimilation and GSH synthesis, or by providing GSH or its precursors directly to the plants. To test if bacterial GSH might be taken up by Arabidopsis to mitigate PEG-induced drought stress, we generated knockout mutants of the microbial GSH biosynthetic genes *gshA* and *gshB*. GSH measurements in both Δ*gshA* and Δ*gshB* mutants revealed a clear reduction in intracellular glutathione levels compared to wild-type SA190 (Figure 5A), with a more pronounced decrease observed in *gshA*. This is consistent with *gshA* catalyzing the rate limiting step in GSH biosynthesis (33). Measurement of oxidized glutathione (GSSG) showed comparable levels in the mutants and wild-type SA190, indicating that the impairment of biosynthesis did not affect GSSG levels (Figure 5B). The GSH/GSSG ratios were significantly lowered in both mutants (Figure 5C), indicating an impaired redox homeostasis and potentially a compromised capacity to detoxify reactive oxygen species (ROS).

**Figure 5.**
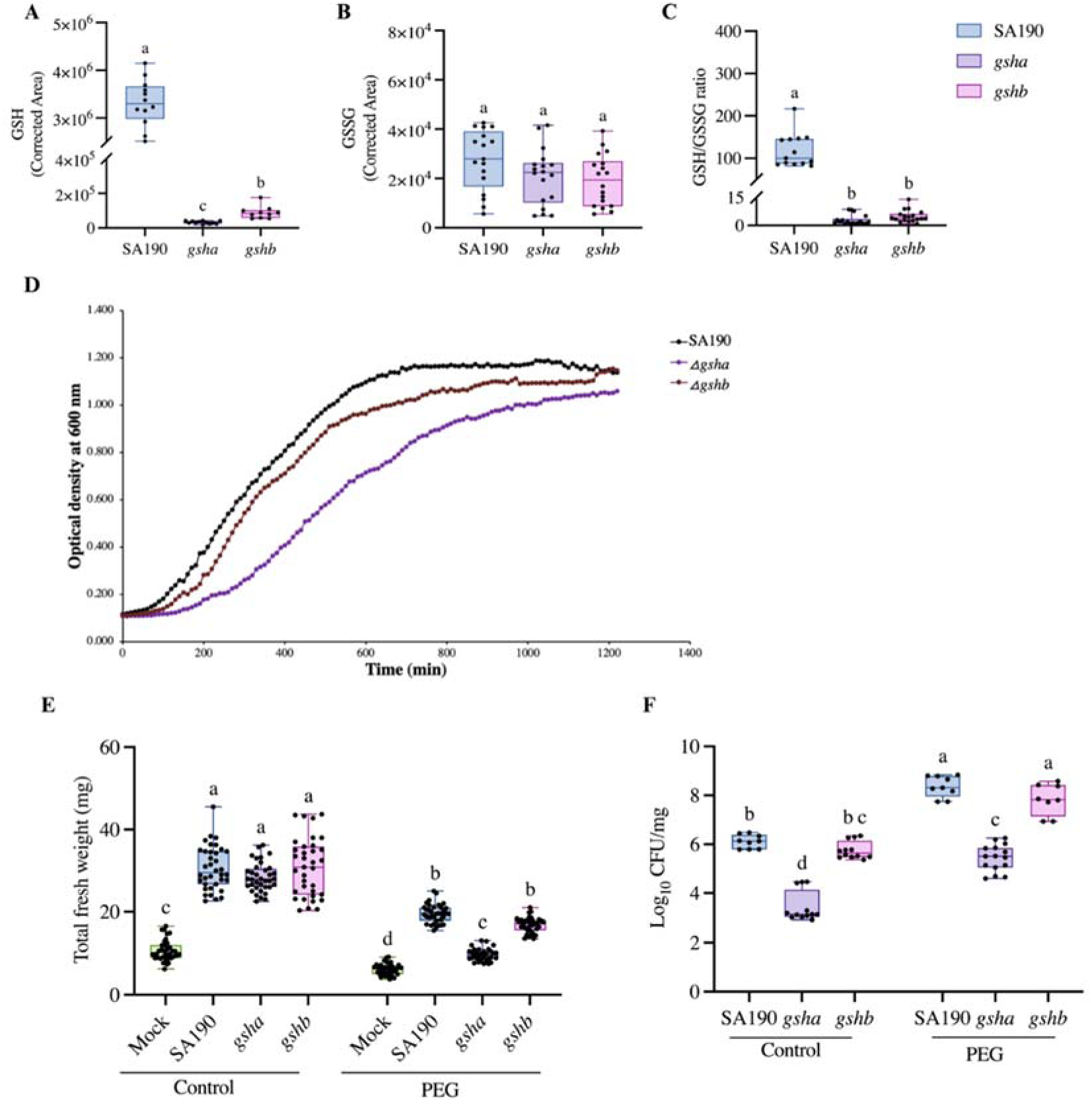
Functional analysis of *Pseudomonas argentinensis* SA190 and glutathione-deficient mutants. Bacterial glutathione (GSH) levels in *SA190 WT*, Δ*gshA*, and Δ*gshB* mutants (A). Bacterial oxidized glutathione (GSSG) levels in *SA190*, Δ*gshA*, and Δ*gshB* (B). GSH/GSSG ratio in *SA190* and the glutathione-deficient mutants (C). For panels A–C, six biological replicates were used. Different letters indicate statistical differences based on one-way ANOVA (p < 0.05). Growth curves of *SA190*, Δ*gshA*, and Δ*gshB* in LB medium (D). Total fresh weight of *Arabidopsis thaliana* Col-0 plants inoculated with *SA190,* Δ*gshA*, or Δ*gshB*, or mock-inoculated, grown under control (non-stress) and 25% PEG-induced drought stress conditions (E). The plant assay was performed with three biological replicates and two technical replicates (n=36). Whiskers indicate minimum and maximum values, the horizontal line marks the median. Statistical differences were determined by two-way ANOVA (*p* < 0.05). Root colonization levels of *SA190* and glutathione-deficient mutants. The boxplots show colony-forming unit (CFU) data from three independent biological replicates, different letters indicate statistical differences based on two-way ANOVA (p < 0.05) (E).

To evaluate the effects of the gene deletions on bacterial fitness, we analyzed the growth curves of both mutant strains as well as SA190 wild-type in LB medium. *gshA* showed a significant growth reduction indicating a vital role of *gshA* in bacterial fitness (Figure 5D). To assess how the deletions of *gshA* and *gshB* affect SA190’s plant growth-promoting ability, we performed plant growth assays under control and PEG-induced drought stress conditions. Under control conditions, seedlings colonized with the Δ*gshA* and Δ*gshB* mutants had biomass levels comparable to seedlings treated with wild-type SA190 (Figure 5E). However, under 25% PEG stress, Δ*gshA* only partially rescued the plant phenotype under PEG-induced drought stress as the plants showed a significantly reduced biomass compared to plants colonized with either SA190 or Δ*gshB*.

Next, plant colonization by the two mutants and wild-type SA190 was compared (Figure 5F). While loss of *gshB* did not affect roots colonization (based on CFU/mg) compared to wild-type SA190, the Δ*gshA* mutant revealed a reduced number of bacteria associated with the root, as indicated by lower CFU/mg values. However, that reduction of bacterial levels does not seem to negatively impact the increased colonization due to PEG-induced stress, as the CFU levels of Δ*gshA* mutant increased (Figure 5F). Together, these findings suggest that *gshA* but not *gshB* is critical for drought stress mitigation effect conferred by SA190.

To further dissect the contribution of bacterial *gshA* and *gshB* and their products to improvement of plant growth under PEG stress, we assessed the interaction of Δ*gshA* and Δ*gshB* mutants with the *cad2-1* mutant. Under control conditions, no obvious differences in phenotypes of *cad2-1* inoculated with the two mutants and wild-type SA190 were observed (Supplementary Figure 8). In contrast, under PEG-induced drought stress, Δ*gshA* was clearly less efficient to rescue the growth inhibition by PEG compared with SA190, whereas Δ*gshB* maintained a phenotype similar to wild-type SA190 (Supplementary Figure 9).

Fresh weight measurements further confirmed these observations. The deletion of *gshA* significantly decreased the ability to increase *cad2-1* biomass under control and PEG stress conditions (Figure 6A and B). In contrast, treatment with Δ*gshB* maintained biomass levels comparable to SA190. Since *cad2-1* has a substantially lower capacity to synthesize GSH, the ability of the Δ*gshB* mutant, which produces γ-glutamylcysteine (γ-EC) but cannot convert it to GSH, to partially restore biomass is notable. This result suggests that γ-EC may be the key bacterial metabolite supporting plant growth under PEG stress. Accordingly, Δ*gshA*, which cannot synthesize γ-EC, failed to enhance growth and biomass accumulation.

**Figure 6.**
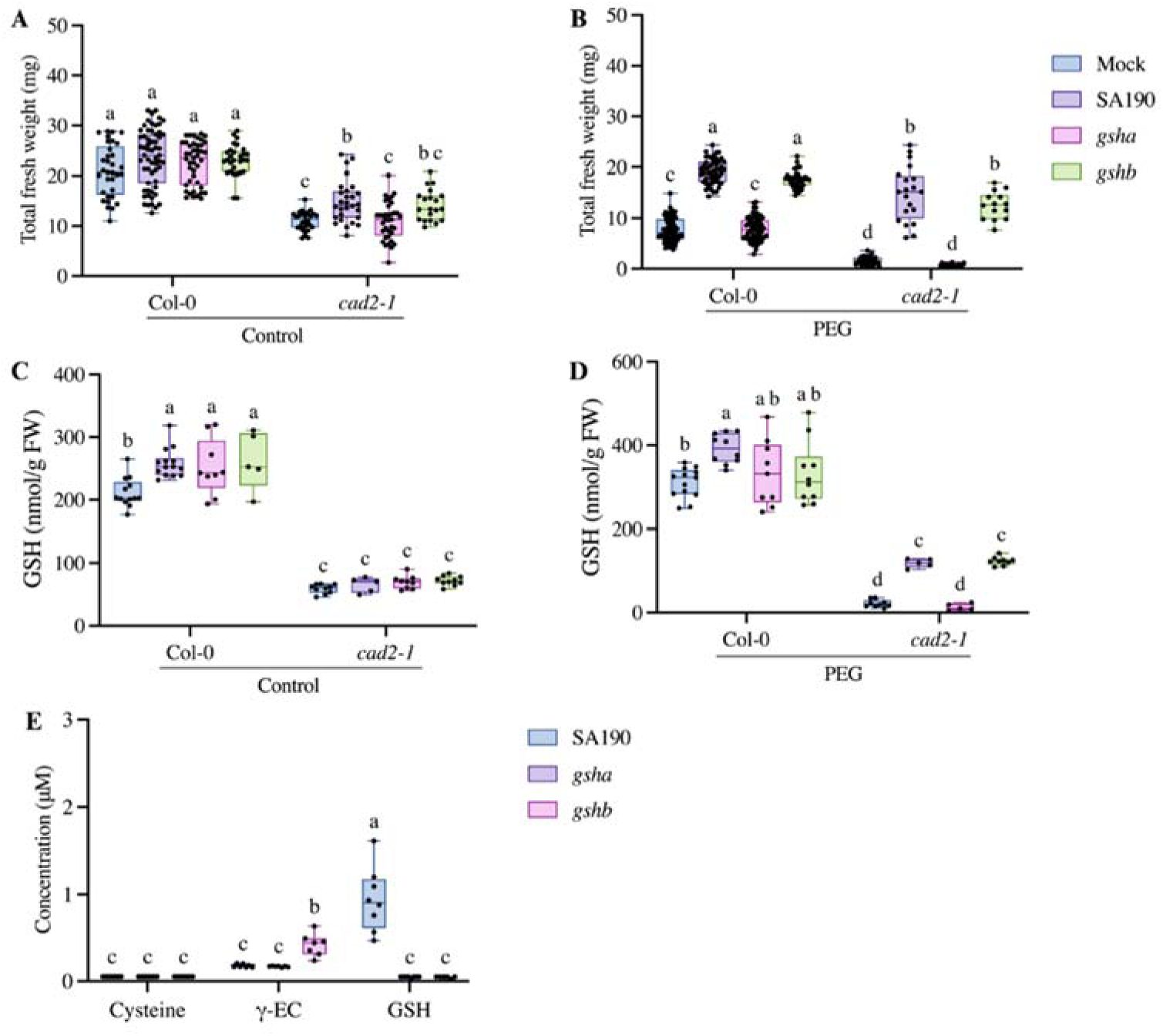
Effects of *Pseudomonas argentinensis* SA190, Δ*gshA*, and Δ*gshB* on *cad2-1* biomass and GSH levels under control and 25% PEG-induced drought stress. Total fresh weight of non-colonized plants, plants colonized with SA190, and plants colonized with Δ*gshA* or Δ*gshB* under control (A) and 25% PEG-induced drought stress (B). Plant assays were performed using three biological replicates with two technical replicates per treatment (n = 36). Boxplot whiskers represent minimum and maximum values, the horizontal line indicates the median. Different letters indicate statistically significant differences based on two-way ANOVA (p < 0.05). Glutathione (GSH) levels in shoot tissue under the same treatments in control (C) and 25% PEG-induced drought stress (D). Different letters indicate statistically significant differences based on two-way ANOVA (p < 0.05). Cysteine, γ-glutamylcysteine (γ-EC), and GSH levels in pellet of SA190 wild type, Δ*gshA*, and Δ*gshB* mutants grown in M9 medium (E). Seven biological replicates were analyzed. Different letters indicate statistically significant differences based on two-way ANOVA (p < 0.05).

GSH quantification from the same samples further supported our conclusion. Under control conditions, inoculation with none of the three strains significantly altered GSH levels in *cad2-1*, while all three promoted GSH accumulation in the wild-type Col-0 (Figure 6C). However, under PEG stress, Δ*gshA* treatment failed to increase GSH accumulation compared with SA190, whereas Δ*gshB* increased GSH levels similar to SA190 (Figure 6D). To verify whether thiols are synthesized in sufficient quantities by the bacteria, we measured cysteine, γ-glutamylcysteine (γ-EC), and GSH levels in wild-type, Δ*gshA*, and Δ*gshB* (Figure 6E). Cysteine levels were comparable in the wild type and the mutants. In contrast, γ-EC levels were significantly elevated in the Δ*gshB* mutant relative to the wild type and Δ*gshA*, consistent with disruption of *gshB* leading to γ-EC accumulation. Both Δ*gshA* and Δ*gshB* mutants exhibited markedly reduced GSH levels compared to the wild type (Figure 6E). These results indicate that SA190-derived γ-EC may contribute to GSH accumulation and biomass enhancement in *cad2-1* under PEG-induced drought stress.

Taken together, these results indicate that bacterial γ-EC and GSH biosynthesis in SA190 is essential for the enhancement of Arabidopsis performance under PEG-induced drought stress. SA190-associated changes in plant sulfur-related metabolites and redox homeostasis further suggest direct contribution of bacterial sulfur metabolites in modulating host responses to PEG stress.

## Discussion

The role of sulfur metabolism in plant adaptation to abiotic stress is increasingly being recognized (34, 35). Numerous studies have demonstrated that plants exhibit an elevated demand for reduced sulfur under stress conditions (35, 36). Reduced sulfur is required for the biosynthesis of cysteine, methionine, and the antioxidant glutathione (GSH), which play central roles in plant defense, redox signaling, and detoxification of reactive oxygen species (ROS) (26).

In this study, we demonstrate that the beneficial endophyte *Pseudomonas argentinensis* SA190 enhances plant drought tolerance through a mechanism involving sulfur metabolism. Measurements of sulfate and GSH levels in plants colonized with SA190 revealed that SA190 significantly increased the abundance of both metabolites in Arabidopsis under PEG-induced drought stress. Consistent with these findings, previous studies have shown that cysteine and GSH levels increased threefold in *Brassica napus* (37), while GSH and ROS detoxifying enzymes, such as glutathione peroxidase, were elevated in *Moringa oleifera* Lam. under salt stress (38). Together, these observations indicate a clear involvement of sulfur-related metabolites in the mitigation of abiotic stress response in plants.

The importance of sulfur metabolism in SA190-mediated stress tolerance was also confirmed that, under PEG-induced drought stress condition, SA190 rescued the severe growth phenotypes of the mutants *sultr1;2*, *apr2*, and *cad2-1*, which are impaired in sulfate uptake, sulfate assimilation, and glutathione biosynthesis, respectively. These data demonstrate that PGPB can influence plant sulfur metabolism, potentially through the supply of sulfur-containing compounds or by modulating the host’s endogenous biosynthetic machinery. Additional experiments revealed that SA190 may directly supply sulfur-containing metabolites to host plant undergoing PEG-induced drought stress. Notably, SA190 treatment markedly increased GSH levels in Arabidopsis exposed to PEG-induced drought stress even when plant GSH biosynthesis was compromised either chemically through BSO or genetically in the *cad2-1* mutant, further supporting this hypothesis.

A growing body of literature indicates that several PGPB improve plant survival under abiotic stress by enhancing sulfur solubility (39), recycling (40), or by modulating the expression of sulfur transport and biosynthesis genes in the host. However, evidence for a direct supply of bacterial sulfur-containing metabolites to plants remains limited. A notable exception is the PGPB strain SA187, which supplies 2-keto-4-methylthiobutyric acid (KMBA) to the plant, leading to increased sulfur content in roots and enhanced survival under salt stress (18).

The elevated GSH levels observed in plants treated with SA190 in this study effectively alleviated oxidative stress and helped maintain cellular redox balance under PEG-induced drought stress conditions. Previous studies have established GSH/GSSG ratios as a reliable indicator of cellular redox status and oxidative stress (41, 42). Measurements of the GSH/GSSG ratios in Arabidopsis colonized with SA190 under PEG-induced drought stress showed a significant increase in reduced GSH relative to the oxidized pool. Consistently, SA190-colonized plants exhibited decreased superoxide accumulation and oxidative damage when exposed to PEG-induced drought stress, further supporting the notion that SA190 may contribute GSH to the plant to maintain redox homeostasis under stress conditions.

SA190 treatment affects plants similarly to exogenous GSH application, which has been shown to alleviate water deficit stress in *Phaseolus vulgaris* plants grown in saline soil (43). Similarly, exogenous GSH application in mung bean (*Vigna radiata* L. cv. Binamoog-1) resulted in enhanced activity of antioxidant and glyoxalase systems and improved GSH/GSSG ratios that were otherwise reduced under PEG-induced drought stress conditions (44). Metabolic profiling further revealed that SA190 treatment significantly increased glutamate and serine levels in Arabidopsis shoots under PEG-induced drought stress. The biosynthetic pathways of these amino acids contribute directly to GSH biosynthesis, thereby linking nitrogen and sulfur metabolism within the plant. These data indicate a coordinated metabolic adjustment mediated by SA190 and suggest an additional route through which SA190 contributes to host metabolic reprogramming by increasing the availability of precursors required for efficient antioxidant production. Taken together, these findings support a model in which SA190 orchestrates a metabolic response that strengthens redox homeostasis in Arabidopsis under PEG-induced drought stress.

Microbial GSH biosynthesis occurs through two ATP-dependent reactions catalysed by the enzymes encoded by the *gshA* and *gshB* genes (45). These genes are essential for maintaining cellular redox balance and are directly implicated in bacterial stress tolerance and virulence (46, 47). In SA190, deletion of *gshA* but not *gshB* inhibited growth, consistent with *gshA* encoding the enzyme responsible for the rate-limiting step of GSH biosynthesis (45). This is similar to plants, where a full knock-out of γ-ECS is embryo lethal while loss of GSH synthetase allows production of viable seeds, although seedlings do not reach maturity (48, 49). Loss of *gshA* and *gshB* not only reduced intracellular GSH levels but also impaired the strain’s ability to alleviate drought stress of inoculated Arabidopsis. These results indicate that SA190-mediated drought tolerance depends on bacterial GSH biosynthesis and supports the hypothesis that SA190 promotes plant stress resilience by reinforcing antioxidant defences.

While all results with the wild type plants or the *sultr1;2* and *apr2* mutants could be explained by SA190 stimulating sulfate uptake and metabolism, the results of the combined analysis of bacterial GSH biosynthesis mutants and the plant *cad2-1* mutant can only be explained by a direct supply of GSH and/or related compounds from the bacteria to the plant host. While neither Δ*gshA* nor Δ*gshB* are disturbed in GSH synthesis, Δ*gshB* is able to produce γ-EC. Since GSH synthetase is intact in *cad2-1* mutant, the PGP effect of Δ*gshB* under PEG-induced stress can be explained by delivery of γ-EC to the plants. Whether γ-EC is the sole metabolite transferred from the bacteria to plants, or whether in wild type situation it is GSH, still needs to be elucidated.

Our findings provide evidence for a potential supply of microbial sulfur metabolites to host plants during plant–microbe interactions. Supporting literature remains limited, although several studies have suggested that soil microbes alleviate drought stress by influencing plant sulfur assimilation or by transforming sulfur into bioavailable forms (50). For example, *Pseudomonas putida* S-313, a strain deficient in sulfonate desulfurization, failed to stimulate tomato growth compared with the wild-type strain (50). Similarly, the Δ*gshA* mutant of *Pseudomonas* sp. 6A2 exhibited reduced biomass under sulfur-deficient conditions (51). Importantly, this study demonstrated that extracellularly secreted GSH from rhizosphere bacteria enhanced plant tolerance to sulfur deficiency (51). Together, these findings underscore a conserved link between microbial sulfur metabolism and plant growth promotion.

In conclusion, our results demonstrate that SA190’s ability to synthesize GSH is critical not only for its own survival and colonization but also for promoting plant drought tolerance. These findings highlight a microbial contribution to plant sulfur metabolism and redox homeostasis during stress. Future studies should investigate whether similar mechanisms operate in crop species under drought conditions. A deeper understanding of plant–microbe interactions in sulfur metabolism may ultimately facilitate the development of innovative strategies to engineer stress-tolerant crops.

## Materials and Methods

### Bacterial Strains and Growth Conditions

*Pseudomonas argentinensis* SA190 (17), *ΔgshA*, and *ΔgshB* mutants were grown overnight at 28 °C on Luria-Bertani (LB, Invitrogen) supplemented with 0.9% agar (Sigma-Aldrich).

### Plant Assays

Plant assays were conducted as previously described by Alwutayd et., 2023 (18) with minor modifications. *Arabidopsis thaliana* ecotype Columbia-0 (Col-0) and the mutant lines *sultr1;2* (19–21), *cad2-1* (22), and *apr2* (*23*) were used in this study. Seeds were surface sterilized with 70% ethanol + 0.05% SDS for 10 min with shaking at 3000 rpm. The seeds were washed three times with 1 mL of 100% ethanol. SA190 and mutants were streaked onto LB plates. Plates were incubated overnight at 28 °C. A single colony from the plates was transferred to LB broth medium. Liquid cultures were incubated overnight at 28 °C, 220 rpm. 100 μL was taken from the overnight culture and transferred to fresh LB broth medium. Incubation at 28 °C was followed by centrifugation at 3000 rpm for 15 min. The bacterial culture was washed with ½ MS solution (Murashige and Skoog basal salts, Sigma-Aldrich). The optical density of the bacterial suspension was set to OD_600_ 0.21. 100 μL of bacterial suspension was added to 50 mL of ½ MS agar and poured into the square Petri plate (120 mm x 120 mm). 100 μL of ½ MS solution was added to 50 mL of ½ MS agar for the mock treatment. Sterilized seeds were distributed on the plates. The plates were covered with aluminum foil and kept in the dark at 4 °C for two days. They were transferred to the growth chamber under long-day photoperiod conditions (16 h light, 8 h dark). After five days of germination, seedlings with 1.0–1.5 cm root length were transferred to ½ MS (non-stress) and ½ MS + 25% PEG plates. Six seedlings were transferred for each plate. They were placed into the growth chamber under the same conditions. The fresh weight of the seedlings was measured 15 days after the seedlings were transferred to ½ MS (non-stress) and ½ MS + 25% PEG plates (PEG-induced drought stress) plates. Primary root length was measured using ImageJ software (Version 1.53t). Root colonization levels were quantified according to the method described by Saad et al. (2018) (52).

### Buthionine Sulfoximine (BSO) Inhibition Assay

Five-day-old non-colonized and SA190-colonized *Arabidopsis thaliana* Col-0 seedlings were transferred to treatment plates containing control, 1 mM buthionine sulfoximine (BSO), or 0.5 mM reduced glutathione (GSH). For the BSO and GSH treatments, stock solutions were added to both the ½ MS agar and the 25% PEG solution to achieve final concentrations of 1 mM BSO and 0.5 mM GSH. As GSH lowers the pH of the media, the pH of both the ½ MS and the PEG solutions was measured after GSH addition and adjusted to 5.7 using sterile 1 M KOH or 1 M HCl. Plates were kept in a growth chamber at 22 °C under long-day conditions (16 h light / 8 h dark). After 12 days of treatment, plants were harvested for fresh weight and thiol quantification.

### Sulfate Measurement

Sulfate quantification was performed as described by Dietzen et al. (2020) (53) with minor modifications. Shoot and root samples (20 mg) were extracted with 1 mL of sterile water, shaken at 4 °C for 1 h, incubated at 95 °C for 15 min, and centrifuged at 4 °C for 45 min. The resulting supernatant was diluted with sterile water and transferred to ion chromatography vials. Standard curves were prepared using 2 mM K_2_SO_4_. Inorganic anions were analyzed using a Dionex ICS-1100 ion chromatography system equipped with a Dionex IonPac AS22 RFIC 4 × 250 mm analytical column (Thermo Scientific). An eluent containing 4.5 mM Na2CO3 and 1.4 mM NaHCO3 was used as the running buffer.

### Quantification of Cysteine and Reduced Glutathione

For the quantification of cysteine and glutathione, 25 mg of shoot and 10 mg of root tissue were extracted in 0.1 M HCl at a 1:10 (w/v) ratio. After centrifugation at 15,000 rpm for 10 min at 4 °C, 60 μL of the supernatant was mixed with 100 μL of 0.25 M CHES-NaOH buffer (pH 9.4) and 35 μL of 10 mM dithiothreitol (DTT), and incubated at room temperature for 40 min. Subsequently, 5 μL of 25 mM monobromobimane (mBBr) was added for thiol derivatization. After 15 min incubation in the dark, 110 μL of 100 mM methanesulfonic acid was added to terminate the reaction. Samples were centrifuged again at 4 °C for 20 min, and 150 μL of the supernatant was transferred to HPLC vials. Standard curves were prepared using serial dilutions (0–100 mM) of 2 mM L-cysteine and 2 mM reduced glutathione. Thiol derivatives were separated by reverse-phase HPLC using a Eurospher 100-3 C18 column (150 × 4 mm, Knauer) with fluorescence detection (excitation: 380 nm; emission: 470 nm). Two buffers were prepared for the analysis: Buffer C (10% [v/v] methanol and 0.25% acetic acid, pH 3.9) and Buffer D (90% [v/v] methanol and 0.25% acetic acid, pH 3.9), at a constant flow rate of 1 mL min^-1^. Peaks corresponding to cysteine and GSH were identified and quantified based on retention times and standard curves.

### Bacterial Growth Assay

Single colonies of SA190, *ΔgshA*, and *ΔgshB* were inoculated into 50 mL LB broth in Erlenmeyer flasks and incubated overnight at 28 °C with shaking at 200 rpm. Overnight cultures were refreshed, and the initial OD_600_ was adjusted to 0.01. Cultures were then transferred to 48-well plates (Greiner Bio-One, Germany), and different concentrations of buthionine sulfoximine (BSO) were tested to assess its effect on SA190’s growth. Final BSO concentrations in the wells were 0.2 mM, 0.5 mM, 1 mM, and 5 mM. Control wells contained no BSO. OD_600_ values were recorded every 10 minutes using a TECAN plate reader (Infinite M200 pro). Growth curves were generated based on time-course OD_600_ measurements.

### Quantification of Reduced and Oxidized Glutathione by HPLC–MS/MS

For plant samples, approximately 80 mg of shoot tissue and 50 mg of root tissue were homogenized using a tissue lyser. One milliliter of extraction solution was added to each sample, followed by shaking at 700 rpm for 30 minutes at 4 °C. For bacterial samples, overnight cultures of SA190, *ΔgshA*, and *ΔgshB* were grown in 50 mL LB broth at 28 °C with shaking at 200 rpm. The following day, 5 mL of each culture was transferred into fresh LB medium and incubated until reaching an optical density at 600 nm (OD_600_) of 0.4. Cultures were centrifuged at 5,000 rpm for 10 minutes at 4 °C, and the supernatants were discarded. Pellets were resuspended in 1 mL of extraction solution and shaken at 700 rpm for 30 minutes at 4 °C. After extraction, all samples were centrifuged at 16,000 × g for 10 minutes at 4 °C. The supernatants were collected and transferred to HPLC vials. Six biological replicates were used for both plant and bacterial samples.

Reduced (GSH) and oxidized (GSSG) glutathione were quantified using high-performance liquid chromatography coupled to tandem mass spectrometry (HPLC-MS/MS) composed by Vanquish UPLC (VF-P20-A Pump, VF-A10-A Sample Module and VH-C10-A Column Compartment) and TSQ Altis Plus Mass Spectrometer, both from Thermo Scientific. The analysis employed a heated electrospray ionization (H-ESI) source in negative ionization mode, with spray voltage set to –2500 V, sheath gas flow of 50 arbitrary units, auxiliary gas at 10, sweep gas at 1, an ion transfer tube temperature of 325 °C, and a vaporizer temperature of 350 °C. Chromatographic separation was carried out on a ACQUITY UPLC BEH Amide Column (2.1 x 150mm with 1.7 um particles) at a constant flow rate of 0.4 mL/min. The mobile phase consisted of two solvents: Water 10 mM Ammonium Formate + 0.15% FA (solvent A) and ACN + 0.05% FA + 2% Ammonium Formate (1 M) (solvent B). The gradient program was as follows: from 0.0 to 12.0 minutes, the elution profile went from 20% to 55% of solvent A; from 12.0 to 15.0 minutes, the flow was kept at 55% of A; from 15.0 to 15.1 minutes, it returned to 20% of A, which was maintained until 18.0 minutes when the run ended. The MS/MS method monitored multiple reaction monitoring (MRM) transitions for both GSH and GSSG over a scan window of 18 minutes. For reduced glutathione (GSH), two transitions were used. GSH1: m/z 308.1 to m/z 162.1 (collision energy of 16.59 V and RF lens voltage at 49 V), and GSH2: m/z 330.1 to m/z 201.1 (collision energy of 17 V and RF lens voltage at 69 V). For oxidized glutathione (GSSG), the following two transitions were monitored. GSSG1: m/z 613.2 to m/z 484.1 (collision energy of 15.36 V and RF lens voltage at 92 V) and GSSG2: m/z 613.2 to m/z 355.1 (collision energy of 21.47 V and RF lens voltage at 92 V) These parameters were optimized to achieve sensitive and specific detection of glutathione species under the described conditions.

### Nitroblue Tetrazolium (NBT) Staining for Superoxide Detection

Seven-day-old non-colonized and SA190-colonized *Arabidopsis thaliana* Col-0 seedlings were used for nitroblue tetrazolium (NBT) staining to detect superoxide accumulation, following the protocol described by Kumar et al. (2014) (54) with minor modifications. Seedlings were placed in 24-well plates and immersed in freshly prepared NBT staining solution. Plates were wrapped in aluminum foil and incubated overnight at room temperature. The following day, the staining solution was removed, and absolute ethanol was added to each well to decolorize the tissues. Plates were incubated in a boiling water bath for 10 minutes. Seedlings were then transferred onto microscope slides and mounted with 60% glycerol. Superoxide accumulation was visualized under a light microscope.

### Metabolite Analysis

Approximately 90 mg of shoot tissue and 40 mg of root tissue were extracted using a solvent mixture of H_2_O:MeOH:CHCl_3_ (1:2.5:1, v/v/v). Samples were vortexed and shaken for 6 minutes at 4 °C, followed by centrifugation for 5 minutes. The resulting supernatant was used for metabolite profiling by gas chromatography–mass spectrometry (GC-MS) using a 7200 GC-QTOF system (Agilent Technologies, Santa Clara, CA, USA), as previously described (55). For quantification, peak areas were normalized to Ribitol used as internal standards.

### Generation of Marker-Free *gshA* and *gshB* Deletion Mutants

Marker-free deletion mutants of *GshA* and *GshB* were generated in SA190 using a two-step homologous recombination approach. For the construction of the deletion, ∼500 bp region up-and downstream of gene of interest were generated by gene synthesis (GenScript, Singapore). The synthesized fragments were cloned into the suicide vector pK18mobsacB (56) via HindIII and XbaI restriction sites. Sequence of the resulting plasmids were verified by sequencing prior to their introduction into SA190 via electroporation. Transformants were selected on LB agar plates supplemented with 50 μg/mL kanamycin. For counterselection of double-crossover recombinants, colonies were plated onto LB agar containing 15% sucrose. Kanamycin sensitive and sucrose resistant clones were screened for the desired gene deletion by colony PCR and Sanger sequencing. Plasmids and primers used in this study are listed in Supplementary Tables 1 and 2.

### Statistical Analysis

All statistical analyses in this study were conducted using GraphPad Prism (Version 10.2.0). The plant assay was performed with three biological replicates and two technical replicates (n = 36). Colony-forming unit (CFU) data were generated from three independent biological replicates. Analysis of total fresh weight, primary root length, sulfate uptake, cysteine, GSH, GSSG, GSH/GSSG and CFU levels was performed using two-way ANOVA with the post-hoc Tukey test. The heatmap were prepared using the MetaboAnalyst program (version 6.0). The plant assay plates were scanned using the EPSON scanner. The primary root length of the plants was measured using the ImageJ program (Version 1.53t). Letters on the graphs indicate statistically significant differences in mean values (p < 0.05). Data were presented in boxplot graphs, where whiskers represent the minimum and maximum values, and the horizontal line indicates the median.

## Author Contributions

M.M.S., S.K., and H.H., supervised the project; B.E., S.K., and H.H. conceptualized the project; B.E., S.K., and H.H. designed experiments; Experimental works were mainly carried out by B.E. with help of R.S.A, C.M., R.J., C.O., and G.M.T.; B.E. analyzed the data and generated figures; M.T. performed the HPLC–MS/MS; P.W. performed GC–MS and analyzed metabolite profiles; B.E. and B.H. generated the mutants; B.E. and R.S.A. drafted the manuscript, all authors read, edited, and aproved the final manuscript.

## Acknowledgments

The authors would thank all members of Hirt lab, AG Kopriva, the CDA management team, and the Bioscience Core Labs in KAUST for the technical assistance and their help in many aspects of this work. The authors also thank Prof. Dr. Jörn Kalinowski, Center for Biotechnology (CeBiTec), Bielefeld University, Germany for donating the pK18mobsacB plasmid. The work was funded by KAUST fund BAS/1/1062-01-01 to H.H. as part of the DARWIN21 desert initiative (http://www.darwin21.org/). S.K. and B.E. are supported by the Deutsche Forschungsgemeinschaft (DFG) under Germanýs Excellence Strategy – EXC 2048/1 – project 390686111 and under Priority Programme “2125 Deconstruction and Reconstruction of Plant Microbiota (DECRyPT)”, project 401836049. The authors acknowledge the use of ChatGPT for assistance with grammatical corrections and improved phrasing in this manuscript.

