## supplemental figures and table for "Root endophyte sulfur metabolites enhance redox balance and drought tolerance in Arabidopsis"

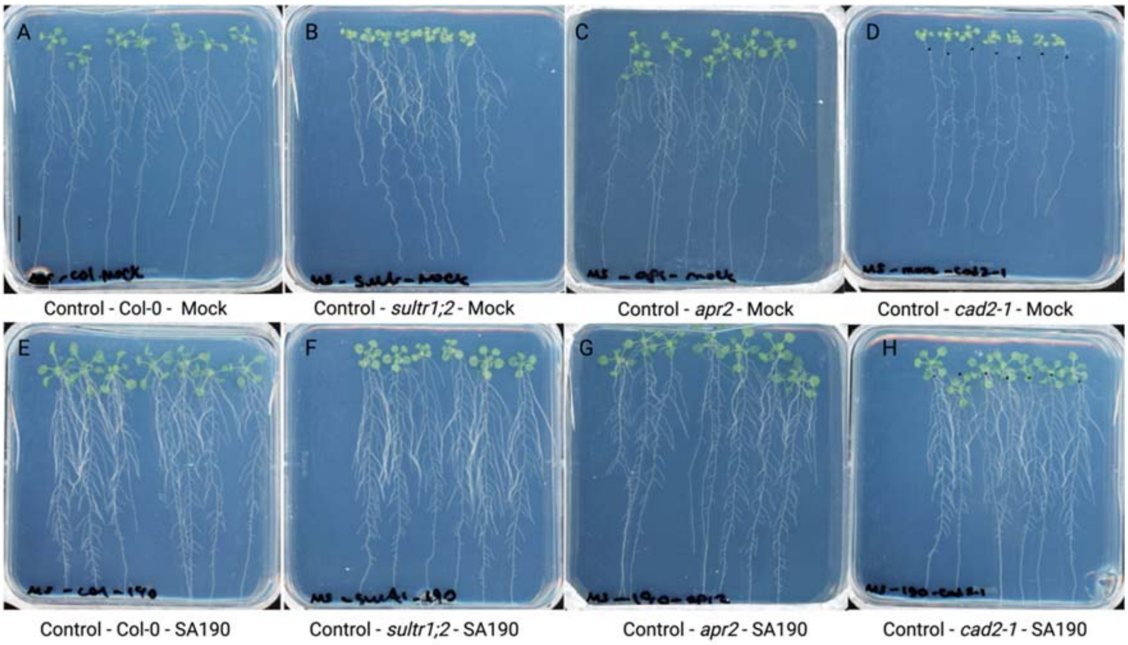


**Fig. S1**. **Effect of *Pseudomonas argentinensis* SA190 on Arabidopsis *sultr1;2, apr2*, and *cad2-1* mutants under control (non-stress) conditions.** Representative phenotypes of *Arabidopsis thaliana* Col-0 seedlings and the mutants after 15 days of growth under control conditions: non-colonized *Arabidopsis thaliana* Col-0, mock (A), non-colonized *sultr1;2,* mock (B), non-colonized *apr2,* mock (C), non-colonized *cad2-1,* mock (D), SA190-colonized *Arabidopsis thaliana* Col-0 (E), SA190-colonized *sultr1;2* (F), SA190-colonized *apr2* (G), SA190-colonized *cad2-1* (H). The scale bar is 1 cm.

.

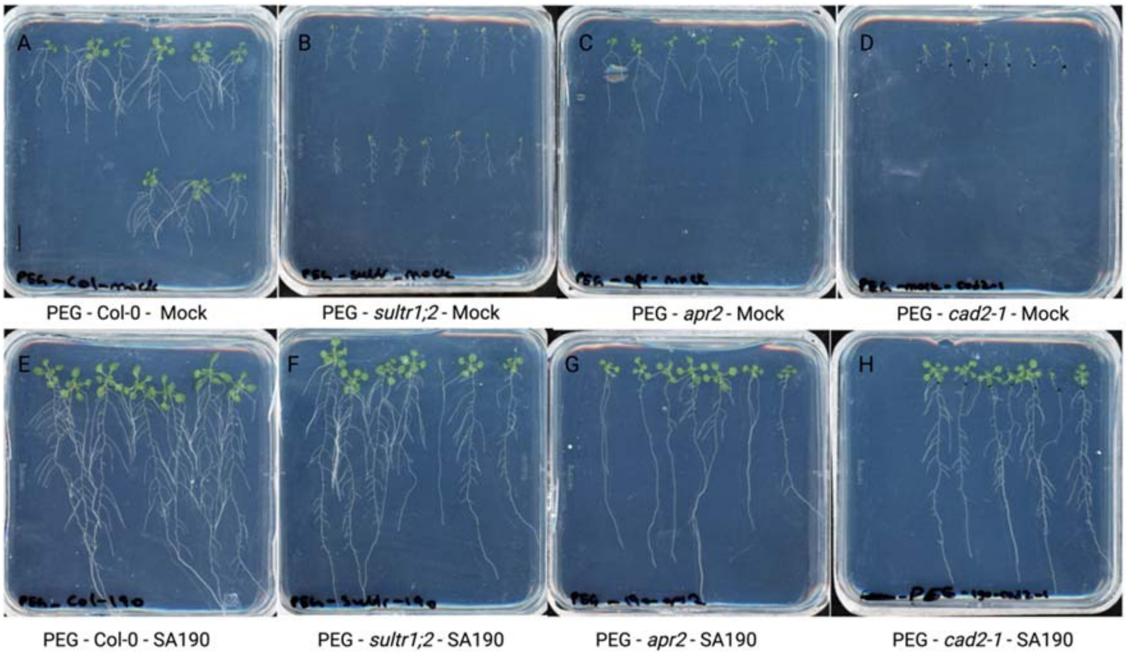


**Fig. S2. Effect of *Pseudomonas argentinensis* SA190 on Arabidopsis *sultr1;2, apr2*, and *cad2-1* mutants under 25% PEG-induced drought stress conditions.** Representative phenotypes of *Arabidopsis thaliana* Col-0 seedlings and the mutants after 15 days of growth under 25% PEG-induced drought stress: non-colonized *Arabidopsis thaliana* Col-0, mock (A), non-colonized *sultr1;2,* mock (B), non-colonized *apr2,* mock (C), non-colonized *cad2-1,* mock (D), SA190-colonized *Arabidopsis thaliana* Col-0 (E), SA190-colonized *sultr1;2* (F), SA190-colonized *apr2* (G), SA190-colonized *cad2-1* (H). The scale bar is 1 cm.

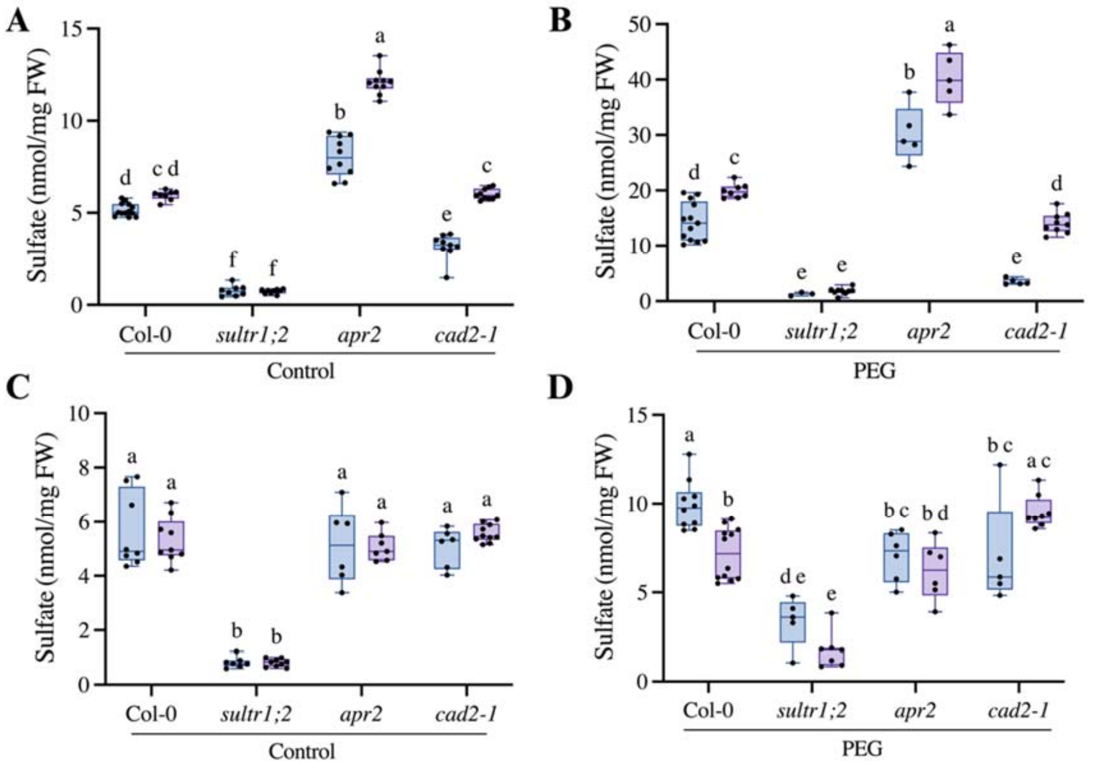


**Fig. S3. Effect of *Pseudomonas argentinensis* SA190 on sulfate uptake in *Arabidopsis thaliana* Col-0 and sulfur pathway-deficient Arabidopsis mutants under control and 25% PEG-induced drought stress conditions.** Sulfate content in shoot samples of Arabidopsis Col-0 and the *sultr1;2, apr2,* and *cad2-1* mutants under control conditions (A) and 25% PEG-induced drought stress conditions (B). Sulfate content in root samples of Arabidopsis Col-0 and the *sultr1;2, apr2,* and *cad2-1* mutants under control conditions (C) and 25% PEG-induced drought stress conditions (D). Whiskers represent minimum and maximum values, the horizontal line indicates the median. Different letters indicate statistically significant differences based on two-way ANOVA ($p < 0.05$).

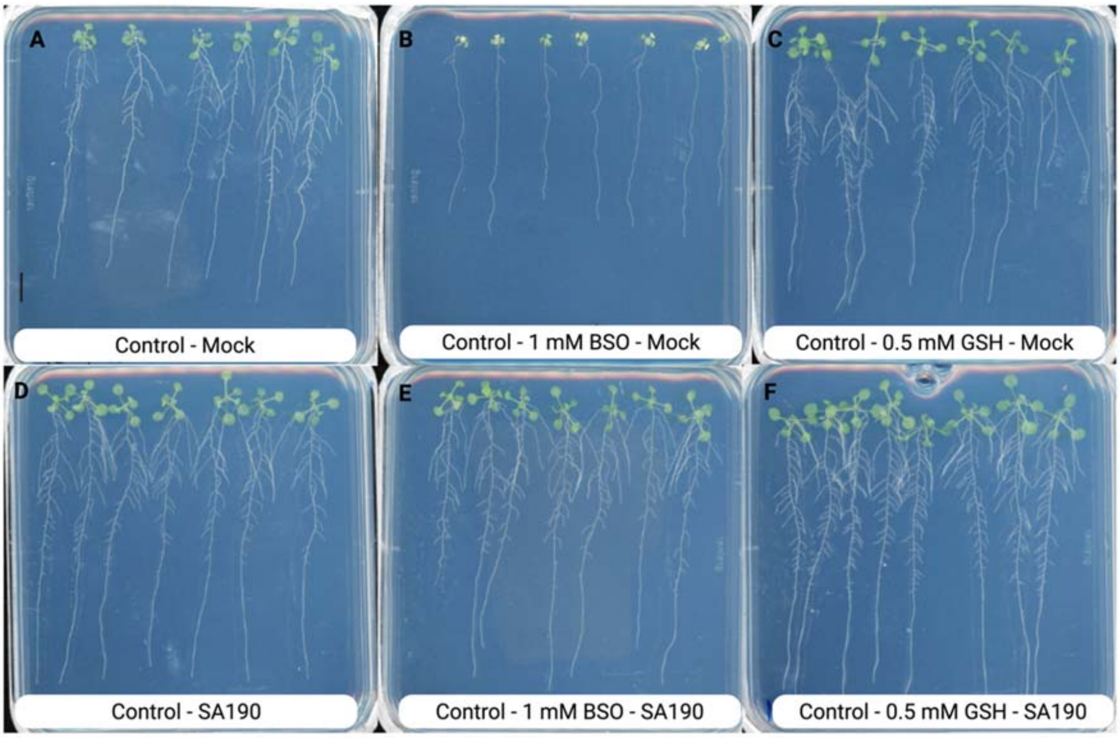


**Fig. S4. Effect of *Pseudomonas argentinensis* SA190 under 1 mM buthionine sulfoximine (BSO) and 0.5 mM glutathione (GSH) treatments in control (non-stress) conditions.** Representative phenotypes of *Arabidopsis thaliana* Col-0 seedlings after 12 days of growth under control conditions: non-colonized control (A), non-colonized + 1 mM BSO (B), non-colonized + 0.5 mM GSH (C), SA190-colonized (D), SA190-colonized + 1 mM BSO (E), and SA190-colonized + 0.5 mM GSH (F). The scale bar is 1 cm.

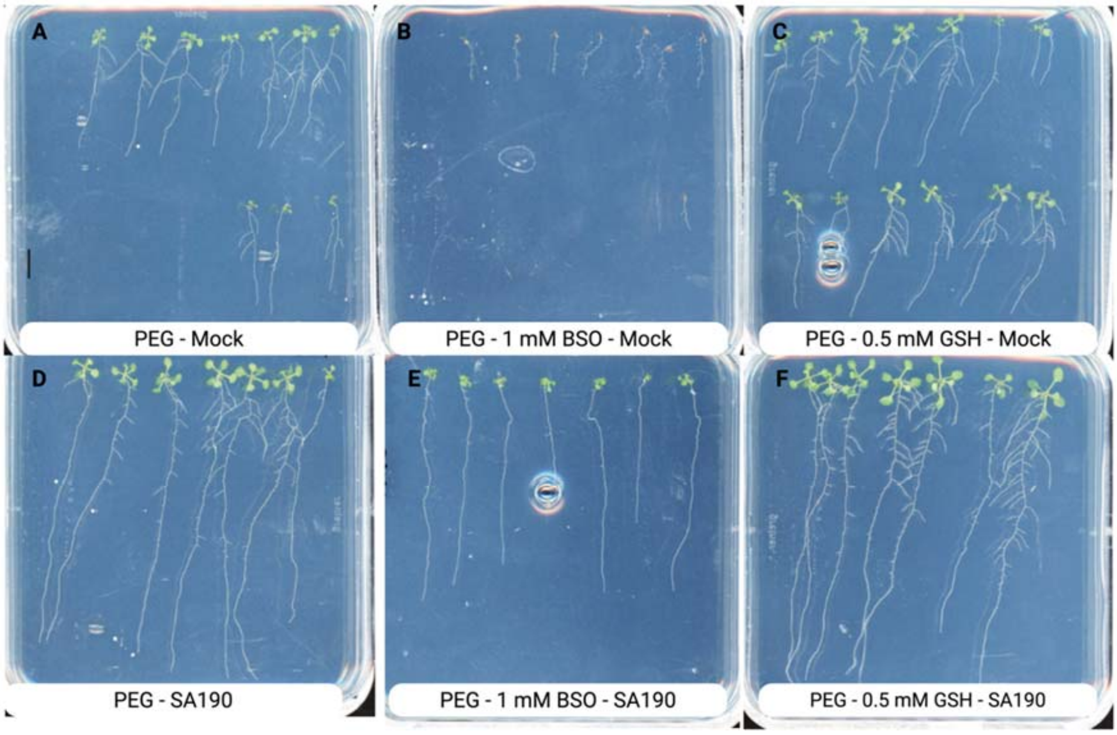


**Fig. S5. Effect of *Pseudomonas argentinensis* SA190 on Arabidopsis under 1 mM buthionine sulfoximine (BSO) and 0.5 mM glutathione (GSH) under 25% PEG-induced drought stress conditions.** Representative phenotypes of *Arabidopsis thaliana* Col-0 seedlings after 12 days of growth under 25% PEG-induced drought stress conditions: non-colonized control (A), non-colonized + 1 mM BSO (B), non-colonized + 0.5 mM GSH (C), SA190-colonized (D), SA190-colonized + 1 mM BSO (E), and SA190-colonized + 0.5 mM GSH (F). The scale bar is 1 cm.

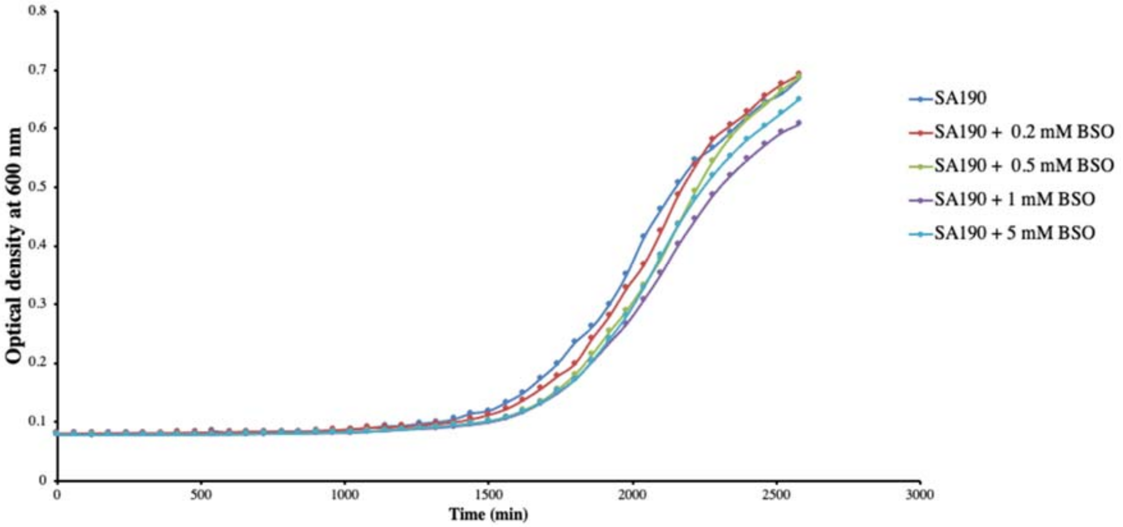


**Fig. S6. Effect of buthionine sulfoximine (BSO) on the growth of *Pseudomonas argentinensis* SA190.** BSO stock solution was added to SA190 liquid cultures to achieve final concentrations of 0.2 mM, 0.5 mM, 1 mM, and 5 mM in M9 medium. Bacterial growth was monitored every 10 minutes using a TECAN plate reader (Infinite M200 pro).

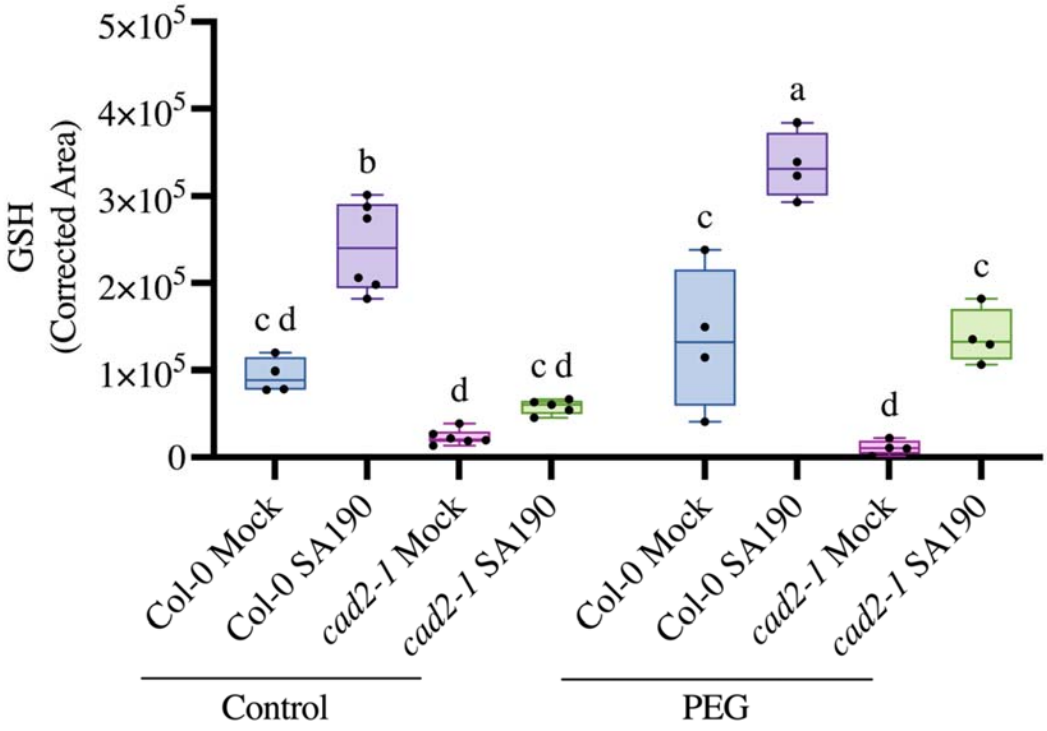


**Fig. S7. Effect of *Pseudomonas argentinensis* SA190 on glutathione (GSH) levels in Arabidopsis *cad2-1* mutant under control (non-stress) and 25% PEG-induced drought stress conditions.** GSH concentrations measured in shoot tissues of *Arabidopsis thaliana cad2-1* seedlings, non-colonized or colonized with SA190, grown under control (non-stress) and 25% PEG-induced drought stress conditions. Boxplots show data from two technical replicates. Whiskers represent minimum and maximum values, the horizontal line indicates the median. Different letters denote statistically significant differences based on two-way ANOVA ($p < 0.05$).

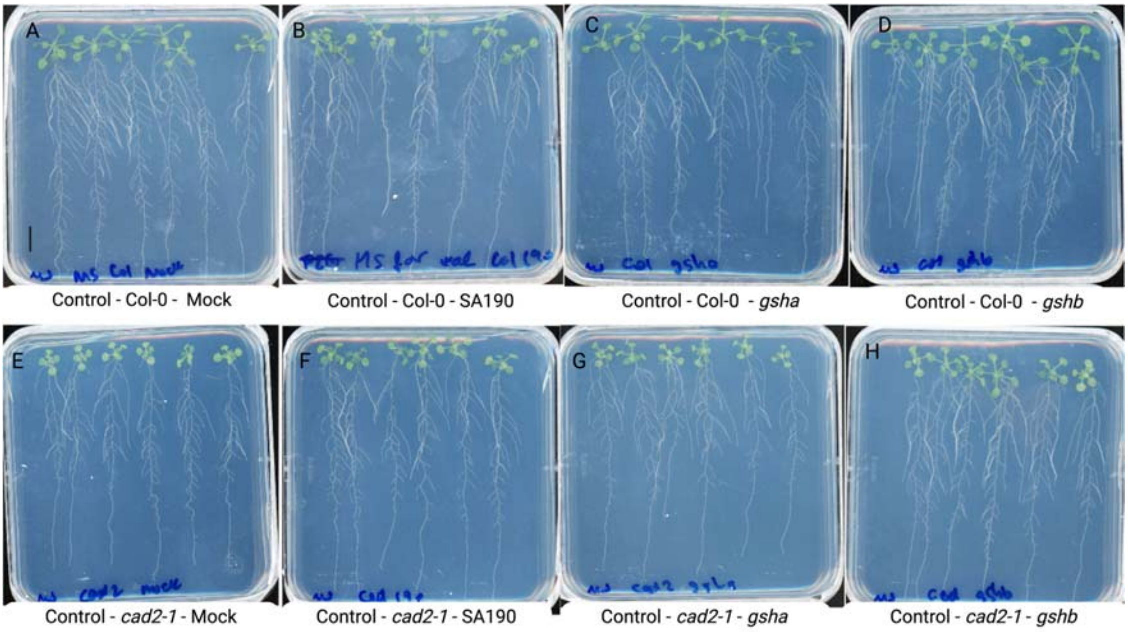


**Fig. S8. Effect of *Pseudomonas argentinensis* SA190 glutathione deficient mutants on Arabidopsis *cad2-1* mutant under control (non-stress) conditions.** Representative phenotypes of *Arabidopsis thaliana* Col-0 and *cad2-1* mutant seedlings after 15 days of growth under control conditions: non-colonized *Arabidopsis thaliana* Col-0, mock (A), SA190-colonized Col-0 (B), Δ*gsha*-colonized Col-0 (C), Δ*gshb*-colonized Col-0 (D), non-colonized *cad2-1*, mock (E), SA190-colonized *cad2-1* (F), Δ*gsha*-colonized *cad2-1* (G), Δ*gshb*-colonized *cad2-1* (H). The scale bar is 1 cm.

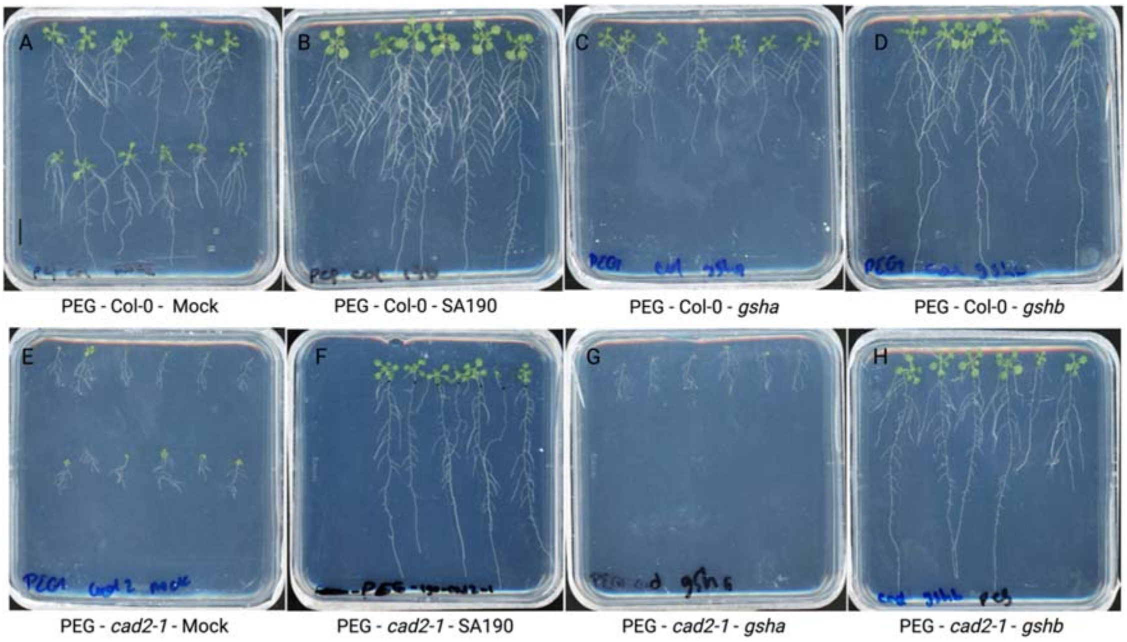


**Fig. S9. Effect of *Pseudomonas argentinensis* SA190 glutathione deficient mutants on Arabidopsis *cad2-1* mutants under 25% PEG-induced drought stress conditions.** Representative phenotypes of *Arabidopsis thaliana* Col-0 and *cad2-1* mutant seedlings after 15 days of growth under PEG stress conditions: non-colonized *Arabidopsis thaliana* Col-0, mock (A), SA190-colonized Col-0 (B), Δ*gsha*-colonized Col-0 (C), Δ*gshb*-colonized Col-0 (D), non-colonized *cad2-1*, mock (E), SA190-colonized *cad2-1* (F), Δ*gsha*-colonized *cad2-1* (G), Δ*gshb*-colonized *cad2-1* (H). The scale bar is 1 cm.

**Table S1.** Plasmids used in this study.

| Plasmids | Characteristics | Reference |
|---|---|---|
| pK18mobsacB | Suicide and narrow-broad-host vector | (Schäfer *et al*., 1994) |

**Table S2.** Primers used for the generation of marker-free *gshA* and *gshB* deletion mutants.

| Primers | Sequence (5'-3') | Usage |
|---|---|---|
| *gsha*_iF | GCCATTCGCCGTCTTTCTTG | |
| *gsha*_iR | CTGCCTGAAGAGGAGGTCAT | |
| *gsha*_oR | GCGACGTGGTCAAGGTCAAG | Deletion of *gsha* |
| *gsha*_oF | GATCACCATGGCGGTGAATT | |
| *gsha*_sF | GTGCTTGCAATCCATCCTAG | |
| *gshb*_iF | GGCACCCTGGTGGTCAAC | |
| *gshb*_iR | CCGCGCATTGAGCTTTTCG | |
| *gshb*_oR | GGACAACTGCGCGCTGTC | Deletion of *gshb* |
| *gshb*_oF | CGGTCGGCGAGTCATCAA | |
| *gshb*_sF | GAAACCAATGGGCCCGGC | |
